# An Automated Bayesian Pipeline for Rapid Analysis of Single-Molecule Binding Data

**DOI:** 10.1101/261917

**Authors:** Carlas S. Smith, Karina Jouravleva, Maximiliaan Huisman, Samson M. Jolly, Phillip D. Zamore, David Grunwald

## Abstract

Single-molecule binding assays enable the study of how molecular machines assemble and function. Current algorithms can identify and locate individual molecules, but require tedious manual validation of each spot. Moreover, no solution for high-throughput analysis of single-molecule binding data exists. Here, we describe an automated pipeline to analyze single-molecule data over a wide range of experimental conditions. We benchmarked the pipeline by measuring the binding properties of the well-studied, DNA-guided DNA endonuclease, TtAgo, an Argonaute protein from the Eubacterium *Thermus thermophilus*. We also used the pipeline to extend our understanding of TtAgo by measuring the protein’s binding kinetics at physiological temperatures and for target DNAs containing multiple, adjacent binding sites.

## Introduction

Single-molecule binding assays allow the interrogation of individual macromolecules from a biological process using purified components or cellular extracts. In contrast to ensemble measurements, single-molecule assays can report the order and kinetics of individual molecular interactions ^1–6^. The introduction of commercial microscopes designed for single-molecule imaging spurred wide adoption of this technology. However, the absence of easy-to-use software with automated pipelines for extracting kinetic data from an image series makes data analysis slow and tedious. Many key steps for obtaining accurate kinetic parameters from co-localization single-molecule spectroscopy (CoSMoS) images still require manual user intervention and the selection of parameters guided by user experience ^7–9^. User-dependent parameter choice and manual inspection of images dramatically limits throughput. For example, after spots are detected via user-defined intensity and bandpass-filter thresholds, the user must still inspect the images to remove overlapping spots and false-positive events. Finally, no standard procedure exists to systematically assess the quality of the analysis. To overcome these hurdles, we constructed a pipeline for rapid processing of CoSMoS images while quantitatively assessing experimental data quality. The process automates experimental calibration and high-confidence spot detection and localization using just minutes of computational time. CoSMoS data processing is controlled through a single graphical user-interface, and the modular interface allows individual functional modules to be adjusted for a wide variety of experiments. The pipeline improves detection of co-localization experiments, data analysis speed, and experimental reproducibility.

## Results

### Pipeline development

**Figure 1** shows the key steps in our pipeline. The package includes detailed installation instructions together with print documentation. The interface comprises a series of tabs, each corresponding to a step in the analysis. The user progresses left to right along, but can readily return to an earlier step, with changes propagating to subsequent steps. The pipeline uses Graphics Processing Unit (GPU) processing to achieve rapid analysis and supports multiple graphics cards.

**Figure 1.**
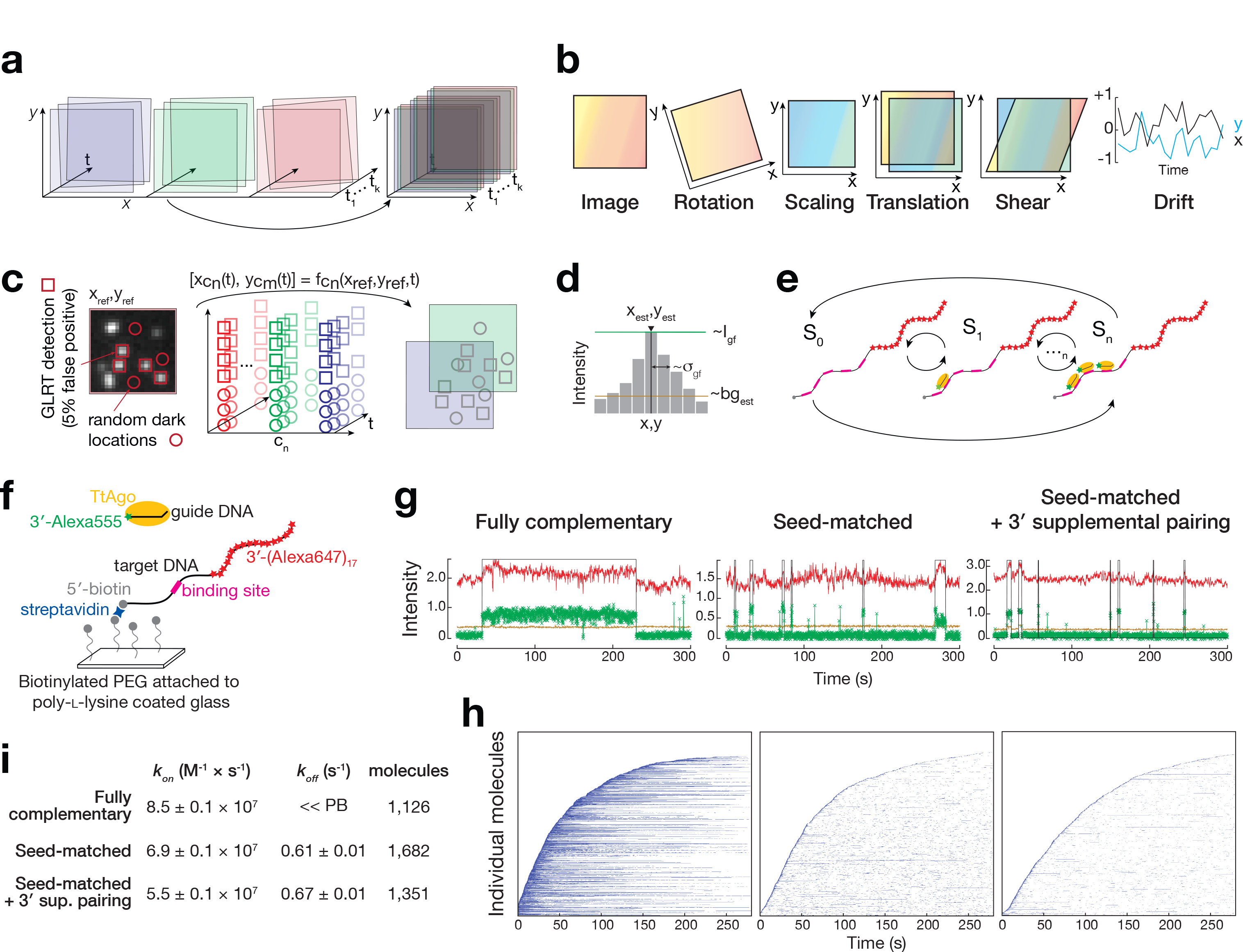
Automated Bayesian Single-Molecule Pipeline for Binding Assays. (**a**) Multiple color channels are registered and corrected for drift. (**b**) Estimated mapping between the colors is time dependent and consists of rotation, scaling, translation, and drift. (**c**) Initial target locations detected by use of Generalized Likelihood Ratio Test (GRLT) are mapped to other views and sub-regions, and are extracted to estimate signal and background parameters. (**d**) Estimated parameters include the position, background, intensity and width of the single-molecule. (**e**) Variational Bayesian Evidence Maximization of Multivariate Gaussian Hidden Markov Model (VBEM-MGHMM) is used to cluster the complexity and estimate parameters of the underlying kinetics. (**f**) Experimental setup to measure TtAgo:guide interactions with target DNA. (**g**) Representative fluorescence intensity time traces of TtAgo (green) binding DNA target (red) with different extents of complementarity to the DNA guide. Light brown indicates background levels of green fluorescence, whereas the black line denotes binding events detected by the pipeline after event filtering (minimal duration and gap closing; Manual – Obtaining the binding traces). This color code is used throughout the Figures in the fluorescence intensity time traces. Fluorescence intensity is expressed in thousands of photons. (**h**) Rastergram summary of traces of individual target molecules, each in a single row and sorted according to their arrival time, for different guide:target pairings. (**i**) Comparison of *k*_*on*_ and *k*_*off*_ of TtAgo with different targets. Values were derived from data collected from several hundred individual DNA target molecules (indicated in the Table as number of molecules); error of bootstrapping is reported.

The first module, *preprocessing*, consists of Electron Multiplying Charge Coupled Device (EMCCD) camera gain calibration, multichannel alignment, and drift correction (**Fig. 1a,b**). The gain and electronic offset of the camera determine the conversion between the number of photons recorded by the camera and the number of digital units contained in the image ^10^. Current CoSMoS methods do not estimate the gain and offset of the cameras, and express signal intensity in arbitrary units. Therefore, parameters required for detection of single molecules are arbitrarily chosen by the user. Because signal-to-background ratios vary between experiments, these parameters should be adapted for every dataset. Based on calibration data, our pipeline estimates gain by exploiting the linear relationship between the noise variance and the mean intensity (see SI Manual – Loading data and gain calibration), allowing automatic parameter estimation and optimal detection, localization and co-localization of single molecules.

After calibrating the gain, fields of view from the wavelength channels corresponding to the different fluorophores used in the experiment must be aligned ^1,7,11^. Alignment corrects differences in rotation, scaling, translation, and shear. The pipeline addresses misalignment by estimating a ‘mapping function’ to relate positions of the target locations in one camera to the mobile components in the other camera. The mapping is obtained via an affine transformation from calibration images of fluorescent beads that emit in both channels (see SI Manual – Alignment of the cameras).

Next, the pipeline corrects for drift caused by movements of the stage ^7,11^. To overcome the need for the traditional fiducial markers, the pipeline estimates drift based on the correlation between consecutive recorded images (see SI Manual – Gain calibration Correction for lateral drift).

The second module, *signal detection and localization*, allows identification of target locations, detection of the binding complexes, and co-localization of the diffusible molecules at each immobilized target (see SI Manual – Target spot detection and Obtaining the binding traces). Current methods identify target positions by using a bandpass filter set by a user-specified intensity threshold ^7,12^. Consequently, considerable manual effort is required to eliminate overlapping spots to prevent the signal from one target molecule from becoming conflated with that from a second, nearby molecule. Unlike methods in current use, the pipeline employs an alternative detection method that uses the photon statistics from the preprocessed images to deliver a minimum number of false-negative detections at a controlled/fixed number of false positives ^13^ (**Fig. 1c**). To automatically eliminate overlapping spots, the pipeline measures the distance from each spot to its neighbors, its circularity, and its width, which enables it to quantitatively discard any spot located within 50 nm of another.

Next, co-localization events are detected. Current methods sum the fluorescence intensity of the mobile component over a small region (~0.4 µm) centered on the mapped and drift-corrected location of the target molecule ^1,2,14^. Co-localization events begin with an abrupt increase and end with an abrupt decrease of the summed fluorescence of the mobile component. To avoid false positives and false negatives, the current methods measure the deviation of the center of mass of the mobile component from the target location ^15^. However, the precision of the position estimation of the center of mass quickly deteriorates with the low signal-to-background ratios often present in CoSMoS experiments ^16^. Thus, abnormally detected events persist and must be removed by visual inspection of the images corresponding to the co-localization intervals, slowing analysis, introducing subjectivity, and degrading reproducibility as noted by Friedman and co-workers ^7^. To address this issue, the pipeline performs maximum-likelihood estimation on the target locations and on the mobile components. This yields an unbiased estimate of the position, local background, spot intensity, and spot width, together with the estimation precision that has the theoretical maximum precision ^17^. Subsequently, these estimates are used by the pipeline to quantitatively score binding events and to define the co-localization intervals. The pipeline requires that authentic binding events meet three user-defined criteria: (1) the mobile component, e.g., an RNA-binding protein, must be detected within a user-specified distance of the target molecule, defined according to the average estimated co-localization precision. The distance between the mobile component and the target location is used to eliminate non-specific binding events caused by protein binding to the cover glass near a target molecule. (2) The spot width must be smaller or equal to the user-specified spot width, defined according to the width of the point-spread function of the microscope ^18^. This criterion ensures that only a single mobile component is specifically bound to the target location. Finally, (3) the fluorescent signal must be above a user-specified signal-to-background ratio, i.e., the fluorescent signal must be a specified number of times greater than the background. This criterion ensures that fluctuations in background fluorescence are not recognized as binding events. This approach also accounts for variations in field illumination, which typically are caused by the relay optics delivering light to the sample ^19^. The pipeline assists the user in setting these criteria by reporting best-practice values for their dataset.

The third module, *data analysis*, calculates association and dissociation rates, as well as the correction for non-specific binding of the mobile component to the glass surface ^7,11^. The data analysis module also estimates the number of complexes bound to target molecules with multiple binding sites (see SI Manual – Analyzing binding kinetics, Correction for the non-specific binding and Hidden Markov Models). Automated analysis of single-molecule data for targets containing multiple binding sites poses a significant technical challenge, because the single-molecule intensity and background fluorescence vary across the field of view. To achieve this, the module uses a Hidden Markov Model (HMM), to determine, based solely on probability, the number of mobile components bound to the same target molecule and the rates of exchange between the different binding states ^20–23^. Multiple HMM analysis frameworks have been proposed to estimate the number of binding states using “information criteria” ^24^. However, when binding events are rare and most target sites are unoccupied, the HMM fit is biased toward an estimate that tries to model the noise due to background fluorescence (also called an unbalanced estimation problem). Furthermore, the number of states of the HMM model is not easy to estimate, because the goodness of the fit increases with additional states. To overcome these two challenges, Bayesian (evidence-based) HMM was introduced by Beal et al. ^25,26^. This approach allows rebalancing the estimation problem using priors to incorporate information known a priori or iteratively estimated. The Bayesian HMM method has been successfully applied to single particle tracking and fluorescence resonance energy transfer, assuming either a zero-mean Gaussian emission distribution ^27^ or a one-dimensional Gaussian emission distribution ^28–30^. Our pipeline extends this framework and enables the estimation of multivariate Gaussians accounting for multi-dimensional, non-zero mean, Gaussian distributed variables ^31^. This permits the use of state estimation in situations where variables are not independent, which is the case for the fluorescence signal and background in CoSMoS experiments.

For each module, all steps are controlled via a user-friendly interface; no knowledge of MatLab syntax or scripting is required. Results from the pipeline can be readily exported to PDF files, and processed data can be exported to MatLab or other software for further analysis. Processed data from an experiment can be saved and merged later with processed data from other replicates in order to estimate the kinetic behavior of the mobile component using a larger number of molecules. Finally, the pipeline uses scripting to save all user-defined parameters, allowing later replication of an experiment or the analysis of another dataset using previously defined parameters.

### Experimental validation of the pipeline

To test the pipeline, we reexamined the binding properties of *Thermus thermophilus* Argonaute (TtAgo), a DNA-guided, DNA-cleaving endonuclease ^32,33^ (**Fig. 1f-i**). TtAgo binds 5′ phosphorylated, 16-nt DNA guides and targets foreign DNA in vivo ^33^. TtAgo pre-organizes the ‘seed’ segment (nucleotides g2–g8) of the guide, pre-paying the entropic penalty for binding the target ^11,32,34–36^. Like other Argonaute proteins, extensive complementarity between the guide and the target allows TtAgo to reach a catalytically competent conformation that can cleave the phosphodiester bond between target nucleotides t10 and t11. Previous single-molecule measurements at 37°C of the on-rate (*k*_*on*_) and off-rate (*k*_*off*_) of TtAgo, guided by a 16-nt DNA corresponding to the first 16 nucleotides of the animal microRNA (miRNA) let-7, revealed that the protein accelerates target finding by >100-times compared to the 16-nt DNA guide in the absence of the protein ^11^. Target complementarity beyond the seed does not increase *k_on_*. TtAgo remains bound to a fully complementary target DNA, but rapidly dissociates from targets complementary to only the seed or the seed plus four 3′ supplementary nucleotides.

Salomon et al. analyzed single-molecule fluorescence images of TtAgo binding ^11^ using imscroll ^7^, a method that identifies co-localization events using high and low intensity thresholds to detect the beginning and the end of a binding event. Because such thresholds cannot be optimal for the entire field of view, Salomon and co-workers manually inspected each binding event analyzed, a process more time consuming than data collection. We compared imscroll to our automated pipeline using single-molecule data for TtAgo:guide DNA complex binding a seed-matched DNA target (**Supplementary Fig. 1**). The pipeline and imscroll detected a similar number of target locations and similar on-(*k*_*on*_^pipeline^ = 7.1 ± 0.1 × 10^7^ M^−1^⋅s^−1^ vs. *k*_*on*_^imscroll^ = 8.6 ± 0.1 × 10^7^ M^−1^⋅s^−1^) and off-(*k*_*off*_^pipeline^ = 0.6 ± 0.01 s^−1^ vs. *k*_*off*_^imscroll^ = 1.0 ± 0.01 s^−1^) rates. Imscroll required 348 of 1,274 putative single target molecules to be manually discarded; the pipeline required no user intervention.

To further test the pipeline, we replicated published experiments analyzing the effect of guide:target complementarity on TtAgo binding ^11^. Using the pipeline to analyze the data gave the expected result that complementarity outside of the seed sequence has little effect on on-rate: fully complementary, *k*_*on*_ = 8.5 ± 0.1 × 10^7^ M^−1^⋅s^−1^; seed only, *k*_*on*_ = 6.9 ± 0.1 × 10^7^ M^−1^⋅s^−1^; seed plus four, 3′ supplementary nucleotides (guide nucleotides g13–g16), *k*_*on*_ = 5.5 ± 0.1 × 10^7^ M^−1^⋅s^−1^. As expected, binding of TtAgo:guide complex to the fully complementary target was too long-lived to permit its off-rate to be measured, because photobleaching of the guide occurred before dissociation. When the target was complementary to just seed or to the seed plus four, 3′ supplementary nucleotides, TtAgo dissociated with the similar, rapid kinetics reported previously (seed only, *τ*_*off*_ = 1.6 s vs. seed plus 3′ supplementary, *τ*_*off*_ = 1.5 s after binding). Thus, our automated approach, using a different method to detect TtAgo binding, calculated *k*_*on*_ and *k*_*off*_ values in good agreement with published results ^11^.

### The pipeline reveals temperature-dependent TtAgo binding dynamics

Previous single-molecule studies examined the binding of the TtAgo:guide complex to DNA and RNA targets at 23°C ^37^, 37°C ^11^, or 45°C ^38^, but *T. thermophilus* grows at 62°C to 75°C ^39^. Thus, knowing the effect of temperature on TtAgo binding is central to understanding the function of the protein in vivo, we measured the temperature dependence of binding kinetics of TtAgo for 285-nt DNA targets with different extents of complementarity to the DNA guide (**Fig. 2**, **Supplementary Fig. 2**). Key to conducting these experiments was our development of an optically transparent sample heater that enables single-molecule experiments at temperatures as high as 55°C. At all temperatures tested, the TtAgo:guide complex bound the three targets with similar, near diffusion-limited on-rates (**Fig. 2a**). Interestingly, mouse AGO2 RISC, which has a similar structure to the TtAgo:guide complex and also possesses endonuclease activity, finds seed-matched targets ~10 times more slowly than fully complementary targets ^11^. Our data suggest that TtAgo does not discriminate between seed-matched and fully complementary targets during its initial search.

**Figure 2.**
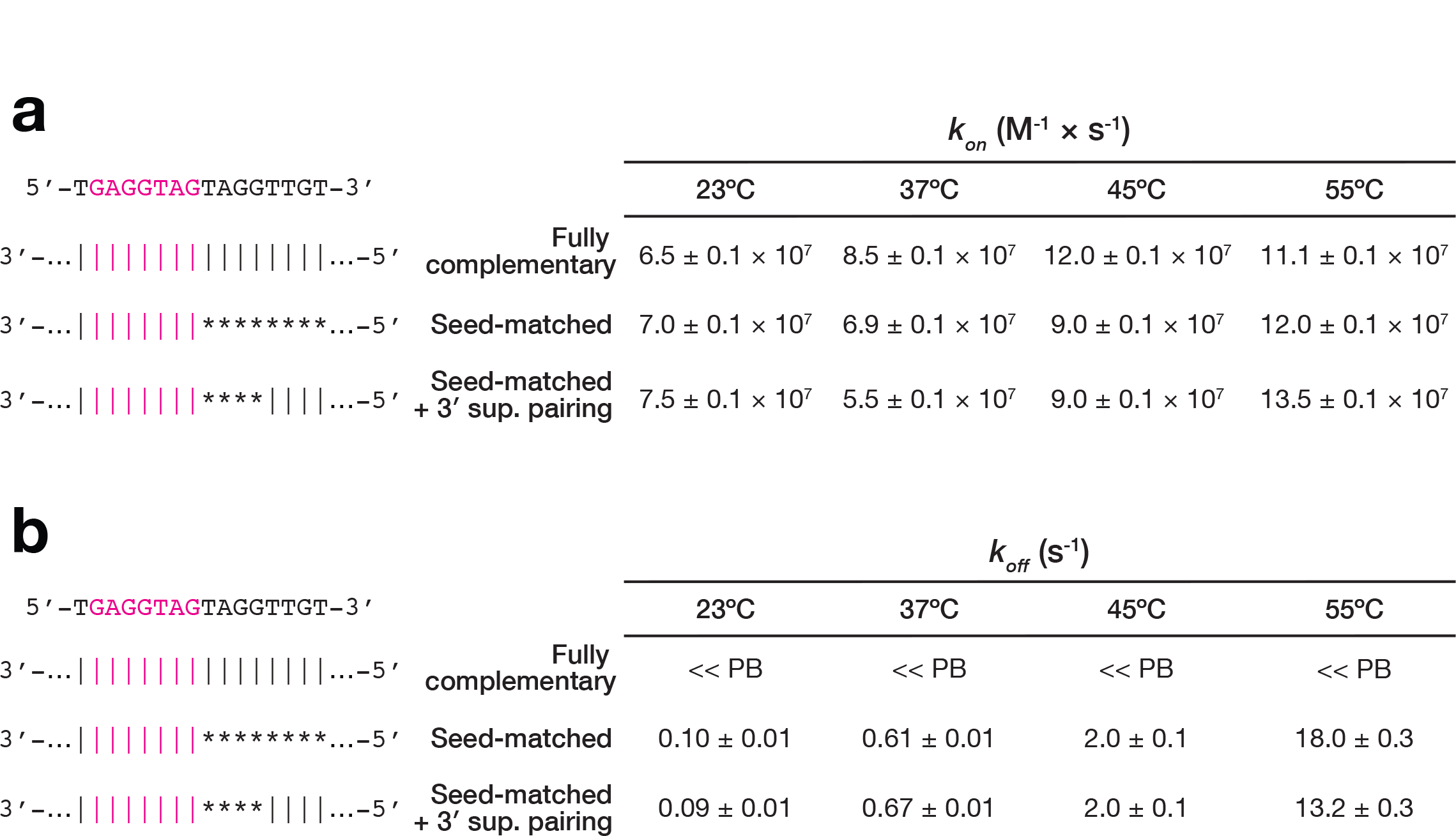
Properties of DNA-guided TtAgo Binding to DNA Targets with Different Extents of Complementarity to the Guide-Strand at Various Temperatures. Values of *k*_*on*_ and **a** *k*_*off*_ (**b**) were derived from data collected from several hundred individual DNA target molecules (>1100); error of bootstrapping is reported.

The dwell time of TtAgo on a target with complete complementarity to the guide remained long and was limited by photobleaching at all temperatures tested. Although at room temperature the TtAgo:guide complex dissociated from targets complementary to the seed or to the seed plus four, 3′ supplementary nucleotides, faster than from the fully complementary target, binding events were stable, τ_*off*_ ~10 s (*k*_*off*_ ~0.1 s^−1^; **Fig. 2b**). Thus, at low temperature, TtAgo displays miRNA-like binding behavior and acts like the RNA-binding, miRNA-guided mammalian Ago2 ^4,11,40^. However, at higher, more physiological temperatures, TtAgo displayed shorter dwell times on targets complementary to the seed or the seed plus four, 3′ supplementary nucleotides, averaging 56 ms (*k*_*off*_ = 18.0 s^−1^) and 76 ms (*k*_*off*_ = 13.2 s^−1^), respectively. Unlike mammalian Ago2, at near-physiological temperature TtAgo binds only transiently to seed-matched targets and requires extensive complementarity to its targets for stable binding. Our data are consistent with the idea that the primary function of TtAgo is to catalyze cleavage of DNA with extensive complementarity to its DNA guide ^34^. The finding that temperature alone, absent any change in amino acid sequence, can convert an Argonaute protein with miRNA-like binding properties into one requiring extensive target complementarity for stable binding, has important implications for the evolution of Argonaute function.

### The pipeline reveals non-cooperative binding of TtAgo to adjacent target sites

In mammals, Argonaute proteins can function cooperatively over short distances, although it is not known whether functional cooperativity reflects cooperative binding ^41,42^. We developed a method based on Variational Bayesian Evidence Maximization (VBEM) and Multivariate Gaussian Hidden Markov Models (MGHMM) to study binding to multiple sites on a single target without the use of additional dyes. To test our method, we performed multi-state analysis of TtAgo binding to DNA targets containing one, two or three binding site(s) fully complementary to the DNA guide. We could detect several TtAgo:guide complexes simultaneously bound to a target molecule, and the pipeline successfully identified the expected number of states (**Supplementary Fig. 3**).

**Figure 3.**
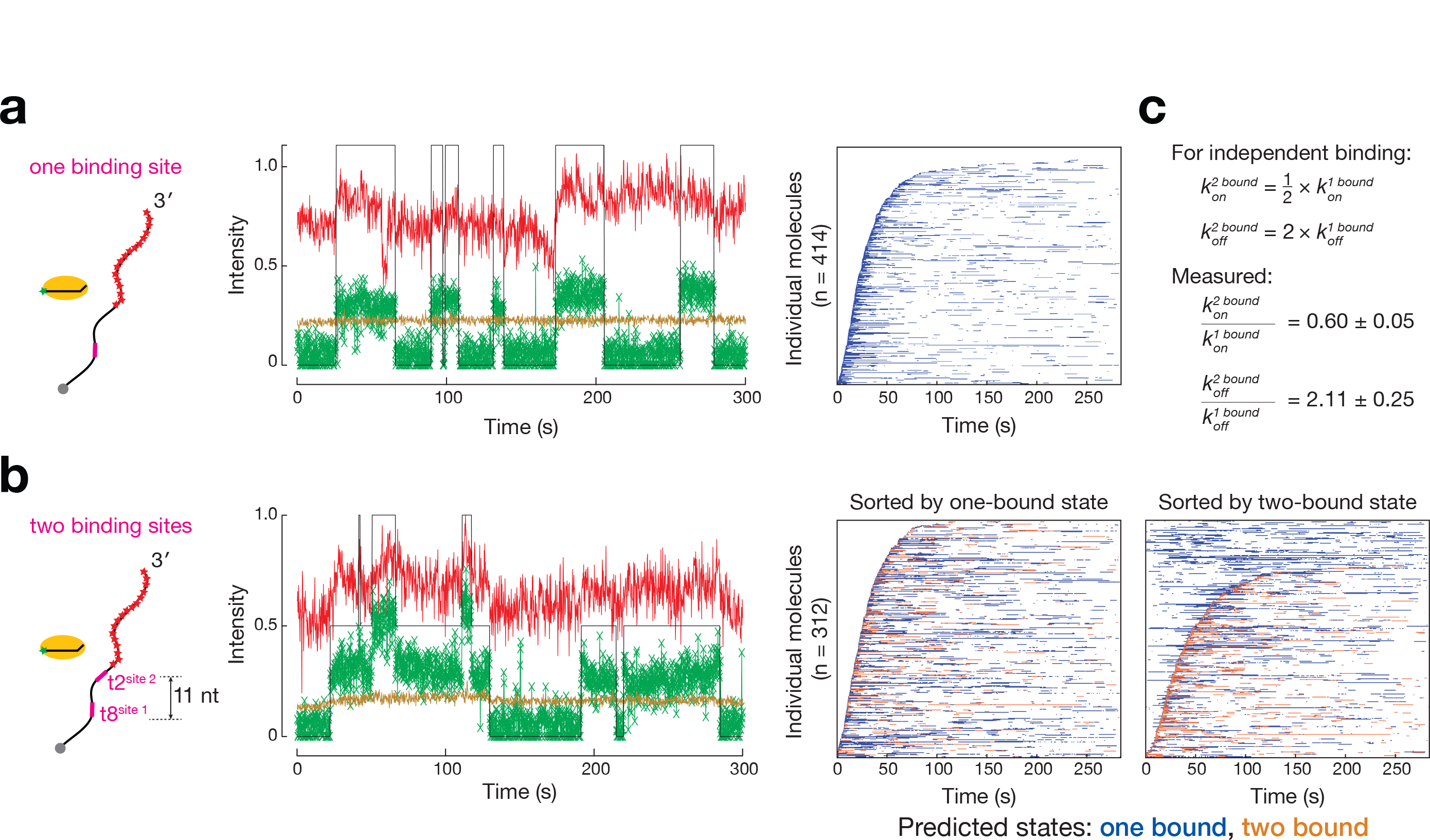
DNA-guided TtAgo Binds Independently to DNA Targets Containing Two Adjacent Seed-Matched t1G Sites. Representative fluorescence intensity time traces of TtAgo (green) binding DNA target (red) containing one binding site (**a**) or two binding sites spaced 11 nt apart from t8 to t2 (**b**). Light brown indicates background levels of green fluorescence, whereas the black line denotes binding events detected by the pipeline after VBEM-MGHMM analysis. Fluorescence intensity is expressed in thousands of photons. Representative rastergrams summarize traces of individual target molecules, each in a single row and sorted according to their arrival time. (**c**) Comparison of *k*_*on*_ and *k*_*off*_ of DNA-guided TtAgo with targets containing one or two binding site(s). Values are reported as mean ± standard deviation for three independent replicates.

Cooperative binding of a complex to one site can either accelerate binding of a second complex at an adjacent site (increasing *k*_*on*_) and/or can stabilize binding at adjacent sites (decreasing *k*_*off*_). To detect differences in binding between multiple and single sites requires a dwell time (1) sufficiently long to allow observation of sequential binding of several TtAgo:guide complexes to the same target molecule, but (2) nonetheless short enough to allow observations to be made before extensive photobleaching occurs. Our standard experimental conditions do not meet these criteria, because TtAgo binding to a seed-matched target is too short to be able to observe simultaneous binding (**Supplementary Fig. 4**), whereas the departure of TtAgo from a fully complementary target is slower than photobleaching (**Fig. 2b**). To circumvent these issues, we used a seed-matched DNA target with deoxyguanosine in the first position (t1G). TtAgo contains a t1G binding pocket ^36,43,44^, and the dwell time of TtAgo for a t1G seed-matched target is >7-times longer (i.e., a smaller *k*_*off*_) than for any other t1N target ^37^ (**Supplementary Fig. 5**). Our DNA guide starts with deoxythymidine (g1T), excluding possible effects of introducing an additional g1:t1 base pair.

Multi-state analysis of TtAgo binding to a DNA target containing two, 7 nt-long, t1G seed-complementary sites 11 nt apart found that *k*_*on*_ for the second site was 0.60 times smaller than for the first site (**Fig. 3**), consistent with a multiple independent sites model (*k*_*on*_^*2 bound*^ = 0.5 *k*_*on*_^*1 bound*^). Supporting this interpretation, *k*_*on*_ for TtAgo binding to a DNA target with two t1G seed-matched binding sites separated by 56 nt was not significantly different from the *k*_*on*_ for the adjacent sites (**Supplementary Fig. 6**). Similarly, *k*_*off*_ for the second site was 2.11 times faster than for the first site (**Fig. 3**), and was not significantly different from *k*_*off*_ when the distance between the two sites was increased (**Supplementary Fig. 6**). As for *k*_*on*_, the *k*_*off*_ values agree well with a model of multiple, independent sites in which *k*_*off*_^*2 bound*^ = 2 *k*_*off*_^*1 bound*^.

## Discussion

We have developed an automated pipeline to analyze single-molecule binding experiments. By eliminating the need for user-supervised rejection of data points, the pipeline reduces analysis times from several weeks for a few hundred traces to a few days for thousands of traces. We validated the pipeline by replicating published results for TtAgo binding kinetics and extended these studies to other temperatures. At near-physiological temperature, TtAgo does not discriminate between miRNA-like targets and siRNA-like targets during the initial search for binding sites, but remains stably bound only to fully complementary targets. Finally, the pipeline, using a VBEM-MGHMM strategy, correctly determines the number of binding sites on a target, allowing us to discover that TtAgo binds independently to adjacent sites.

## Acknowledgements

We thank members of the Grunwald and Zamore laboratories for discussions and comments on the manuscript; Victor Serebrov for experimental and analytical advice; Joerg Braun and Amena Arif for feedback on the pipeline. This work was supported in part by National Institutes of Health grant R37GM062862 to P.D.Z and a junior research fellowship through Merton College, United Kingdom, to C.S.S.

## Author contributions

C.S.S., K.J., D.G., and P.D.Z. contributed to the study design. C.S.S. derived theory, designed, and implemented the pipeline. S.M.J. purified TtAgo:guide complex. K.J. collected data and optimized experimental conditions. K.J. and C.S.S. analyzed data. M.H. led the development of the stage heater. C.S.S. and K.J. initiated, and D.G., and P.D.Z. supervised project. K.J., C.S.S., D.G., and P.D.Z. wrote the manuscript. All authors revised and approved the manuscript.

## Methods

### Data acquisition

Alexa647-labeled target DNA was immobilized on a polymer-coated glass surface via biotin-streptavidin interaction. TtAgo was loaded with a 16 nt, 3’ Alexa Fluor 555-labeled single-stranded DNA guide (see SI - Preparation of TtAgo:guide complex). A syringe pump (KD Scientific, Holliston, MA) running in withdrawal mode at 0.15 ml⋅min^−1^ was applied to the flow cell outlet to introduce TtAgo:guide complex (pre-heated to 23°C, 37°C, 45°C or 55°C) supplemented with an oxygen scavenging system ^45,46^ and triplet quenchers ^47^. Continuous acquisition of frames began when the TtAgo:guide solution was introduced. Typically, 1,500–8,000 frames were collected at 5– 67 frames⋅s^−1^.

Imaging was performed on an IX81-ZDC2 zero-drift inverted microscope equipped with a cell^TIRF motorized multicolor TIRF illuminator with 405, 488, 561, and 640 nm 100 mW lasers and a 100×, oil immersion, 1.49 numerical aperture UAPO N TIRF objective with FN = 22 (Olympus, Tokyo, Japan). Alexa555 and Alexa647 molecules were excited with only the 561-nm laser, as the presence of 17 Alexa647 dyes on the target produces sufficient signal at the lower wavelength. Use of a single laser ensured that both dyes were excited within the same focal volume. Fluorescence signals were split with a main dichroic mirror (Olympus OSF-LFQUAD) and triple emission filter (Olympus U-CZ491561639M). The primary image was relayed to two ImagEM X2 EM-CCD cameras (C9100-23B, Hamamatsu Photonics, Hamamatsu, Japan) using a Cairn three-way splitter equipped with a longpass dichroic mirror (T635lpxr-UF2, Chroma) and bandpass filters (Chroma 595/50) in front of the ‘green’ camera. Illumination and acquisition parameters were controlled with cell^TIRF and MetaMorph software (Molecular Devices, Sunnyvale, CA), respectively. The TIRF imaging system was isolated from floor vibrations with a Micro-g laboratory table (Technical Manufacturing Corporation, Peabody, MA).

A digitally-controlled heater (TP-LH, Tokai Hit) maintained objective temperature at 40°C (except when experiments were performed at 23°C). A custom fabricated heating stage was heated to 45°C, 55°C, or 80°C to achieve sample temperatures of 37°C, 45°C, or 55°C, respectively. Temperature on the surface of the cover glass was independently monitored with a Type E, 0.25 mm O.D. thermocouple (Omega Engineering Inc., Sutton, MA) inserted between the top and the bottom cover glasses.

### Data analysis

Images were recorded as uncompressed TIFF files and merged into stacked TIFF files. Images were processed using the pipeline (SI Manual - collection of a complete dataset). First, 100 images of a grid slide and of background were used to estimate the gain of CCD cameras ^13^. Second, 10 images of fluorescent streptavidin-labeled microspheres (Life Technologies F-8780) were used to determine alignment of images from multiple wavelength channels. Third, lateral drift of the surface was determined for each frame using target molecules as immobilized markers. Locations of target molecules were picked in the first frame acquired by performing a Generalized Likelihood Ratio Test in each pixel ^13^. Large clusters of positive pixels where filtered out, but all identified spots were visually inspected, and locations corresponding to multiple target molecules were removed. To obtain binding traces in all frames the identified locations were fitted using Maximum Likelihood Estimation. Co-localization events required that (1) the intensity of TtAgo complex > 150 photons, (2) ratio intensity of the TtAgo:guide complex to the local background > 1, (3) the distance between the target and guide was < 1 pixel, and (4) sigma < 4.6. To exclude short, non-specific events, the minimal event duration was set to 2–5 frames. To overcome short temporary loss of TtAgo fluorescent signal due to blinking of the fluorescent dye, the gap parameter was set to 2–5 frames. Only the first binding event at each target location was used for estimation of arrival time and dwell time, in order to minimize errors caused by occupation of sites by photobleached molecules. The same analysis was automatically performed on “dark” locations, i.e., regions that contained no target molecules; these served as a control for non-specific binding of TtAgo complex to the surface of the cover glass. The analysis was scripted to ensure reproducibility of user settings. The individual experiments were saved, combined, and error evaluated by 1,000-cycle bootstrapping of 90% of the data.

To calculate the number of binding sites, VBEM-MGHMM analysis was first performed with priors manually estimated from fluorescence intensity time traces (See manual – Hidden Markov Models). The starting point of the signal and background priors, *m*, is set to the mean signal and background of a single binding event of TtAgo. The starting point of priors *k*, *v* and *W*^1/2^ for model order selection are set to 10. Subsequently, the estimated prior parameters (*m*, *k*, *v* and *W*^1/2^) are used to automatically segment the traces with a correct model order ^48^.

## Methods (SI)

### Preparation of TtAgo:guide complex

Expression and purification of TtAgo was essentially as described ^32^. Briefly, TtAgo coding sequence was cloned into pET SUMO (Life Technologies) and expressed in *E. coli* BL21-DE3 by inducing at OD_600_ of 0.5 with 0.2 mM isopropyl-β-D-thiogalactoside at 37°C for 8 h. Cells were lysed (micro-fluidizer, Microfluidics, Westwood, MA), and TtAgo purified by HisTrap HP (GE Healthcare) chromatography. The amino terminal six-histidine tag was cleaved from TtAgo using SUMO-protease (Life Technologies), and the protein was further purified by HiTrap SP HP (GE Healthcare) chromatography. Purified TtAgo was dialyzed into storage buffer (20 mM HEPES-KOH, pH 7.4, 250 mM potassium acetate, 3 mM magnesium acetate, 0.1 mM EDTA, 5 mM dithiothreitol, 20% [w/v] glycerol). TtAgo (0.4 µM) was incubated with 1.2 µM 16 nt, synthetic, single-stranded DNA oligonucleotide corresponding to the first 16 nt of let-7a and bearing a 3′ Alexa555 dye (Invitrogen) for 30 min at 75°C in 20 mM HEPES-KOH, pH 7.4, 350 mM potassium acetate, 3 mM magnesium acetate, 0.01% (w/v) Igepal CA-630, 5 mM dithiothreitol, and 20% (w/v) glycerol. Unassembled DNA guide was removed by passing the loading reaction through a Q Sepharose Fast Flow (GE Healthcare) spin column. TtAgo:guide complex concentration was measured by fluorescence with Typhoon FLA-7000 (GE Healthcare) following denaturing polyacrylamide gel electrophoresis. The complex was flash frozen and stored at ‒80˚C.

### Preparation of DNA Targets

Single-stranded DNA targets were generated by annealing synthetic oligonucleotides to a Klenow template oligonucleotide as described ^11^ (Table S1). In a typical labeling procedure, 100 pmol DNA target was mixed with a 1.5-fold molar excess of Klenow template oligonucleotide in 7.5 µl 10 mM HEPES-KOH (pH 7.4), 20 mM sodium chloride and 0.1 mM EDTA. Samples were incubated at 90°C for 5 min in a heat block. Then, the heat block was switched off and allowed to cool to room temperature. Afterwards, the annealed strands (30% of final reaction volume) were added without further purification to a 3′ extension reaction, comprising 1× NEB buffer 2 (New England Biolabs, Ipswich, MA), 1 mM dATP, 1 mM dCTP, 0.12 mM Alexa Fluor 647-aminohexylacrylamido-dUTP (Life Technologies), and 0.2 U/µl Klenow fragment (3′→5′ exo-minus, New England Biolabs) and incubated at 37°C for 1 h. The reaction was quenched with 500 mM (f.c.) ammonium acetate and 20 mM (f.c.) EDTA. A 15-fold molar excess of “trap” oligonucleotide (Table S1) was added to the Klenow template oligonucleotide. The entire reaction was precipitated overnight at ‒20°C with three volumes of ethanol. The labeled target was recovered by centrifugation, dried, dissolved in loading buffer (7M Urea, 25 mM EDTA), and incubated at 95°C for 5 min. The samples were resolved on 6% polyacrylamide gel and isolated by electroelution.

### Microscope Slide Preparation

Microfluidic chambers were prepared on cover glasses as described ^11^. Briefly, cover glasses (Gold Seal 24 Å~ 60 mm, No. 1.5, Cat. #3423), and glass coverslips (Gold Seal 25 Å~ 25 mm, No. 1, Cat. #3307) were cleaned by sonicating for 30 min in NanoStrip (KMG Chemicals, Houston, TX), followed by washing with 10 changes of deionized water and stored in deionized water. Fresh cover glasses were prepared for each day of imaging. Cover glasses and coverslips were dried with a stream of nitrogen. Two ~1 mm diameter lines of high vacuum grease (Dow Corning, Midland, MI) were applied to the cover glass to create a flow cell. Three layers of adhesive tape were applied outside of the flow cell. The coverslip was placed on top of the cover glass, with a ~0.3 mm gap between the cover glass and coverslip. To minimize non-specific binding of protein and DNA molecules to the glass surface, microfluidic chambers were incubated with 2 mg/ml poly-L-lysine-graft-PEG-biotin in 10 mM HEPES-KOH, pH 7.4 at room temperature for 30 min and washed extensively with LSE buffer (30 mM HEPES-KOH pH 7.9, 120 mM potassium acetate, 3.5 mM magnesium acetate, 20% [w/v] glycerol) immediately before use. To allow immobilization of biotinylated protein or DNA targets, streptavidin (0.01 mg/ml, Sigma) was incubated for 5 min in each chamber. Unbound streptavidin was washed away with LSE buffer.

### Single-Molecule Experiments

The enzymatic oxygen scavenging system comprised 2.5 mM protocatechuic acid (PCA, Aldrich 37580) and 0.5 U/ml *Pseudomonas sp.* protocatechuate 3,4-Dioxygenase (PCD, Sigma P8279). Triplet quenchers, trolox (Aldrich 238813), propyl gallate (Sigma P3130), and 4-nitrobenzyl alcohol (Aldrich N12821) were each added to 1 mM (final concentration).

Immediately before each experiment, a flow cell was incubated with LSE buffer supplemented with 75 µg/ml heparin (Sigma H4784), oxygen scavenging system and triplet quenchers for 2 min. Then, it was filled with ~100 pM target in LSE buffer supplemented with 75 µg/ml heparin, oxygen scavenging system and triplet quenchers. Target deposition was monitored by taking a series of images; once the desired density was achieved, the flow cell was washed three times with LSE buffer supplemented with oxygen scavenging system and triplet quenchers.

---

**Figure S1.**
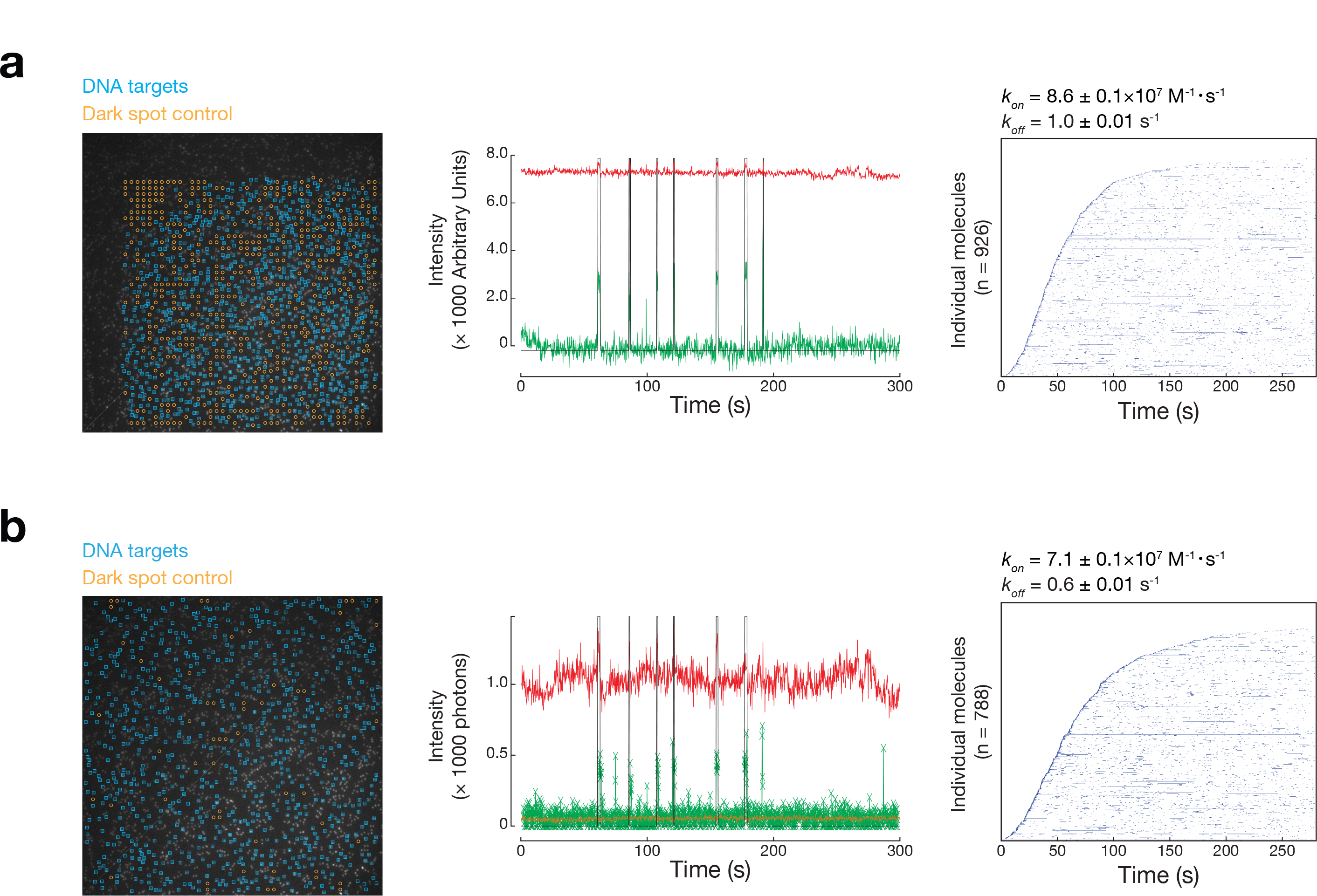
Comparison of the Pipeline to Imscroll, Related to Figure 1. The same data set of TtAgo:guide complex binding a seed-matched target was analyzed by imscroll (**a**) and by the pipeline (**b**). Image representing the selection (blue squares around molecules) of the DNA targets used to analyze DNA-guided TtAgo binding. “Dark” locations, i.e., regions that contained no target molecules (yellow circles) served as a control for non-specific binding of TtAgo:guide complex to the surface of the cover glass. Representative fluorescence intensity time traces obtained by imscroll (**a**) or the pipeline (**b**) for DNA-guided TtAgo (green) binding the same DNA target molecule (red). The black line denotes detected binding events. Light brown indicates background levels of green fluorescence calculated by the pipeline. Imscroll does not provide this information. Rastergrams summarize traces of individual target molecules, each in a single row and sorted according to their arrival time. Values of *k*_*on*_ and *k*_*off*_ were derived from several hundred individual DNA target molecules; error of bootstrapping is reported.

**Figure S2.**
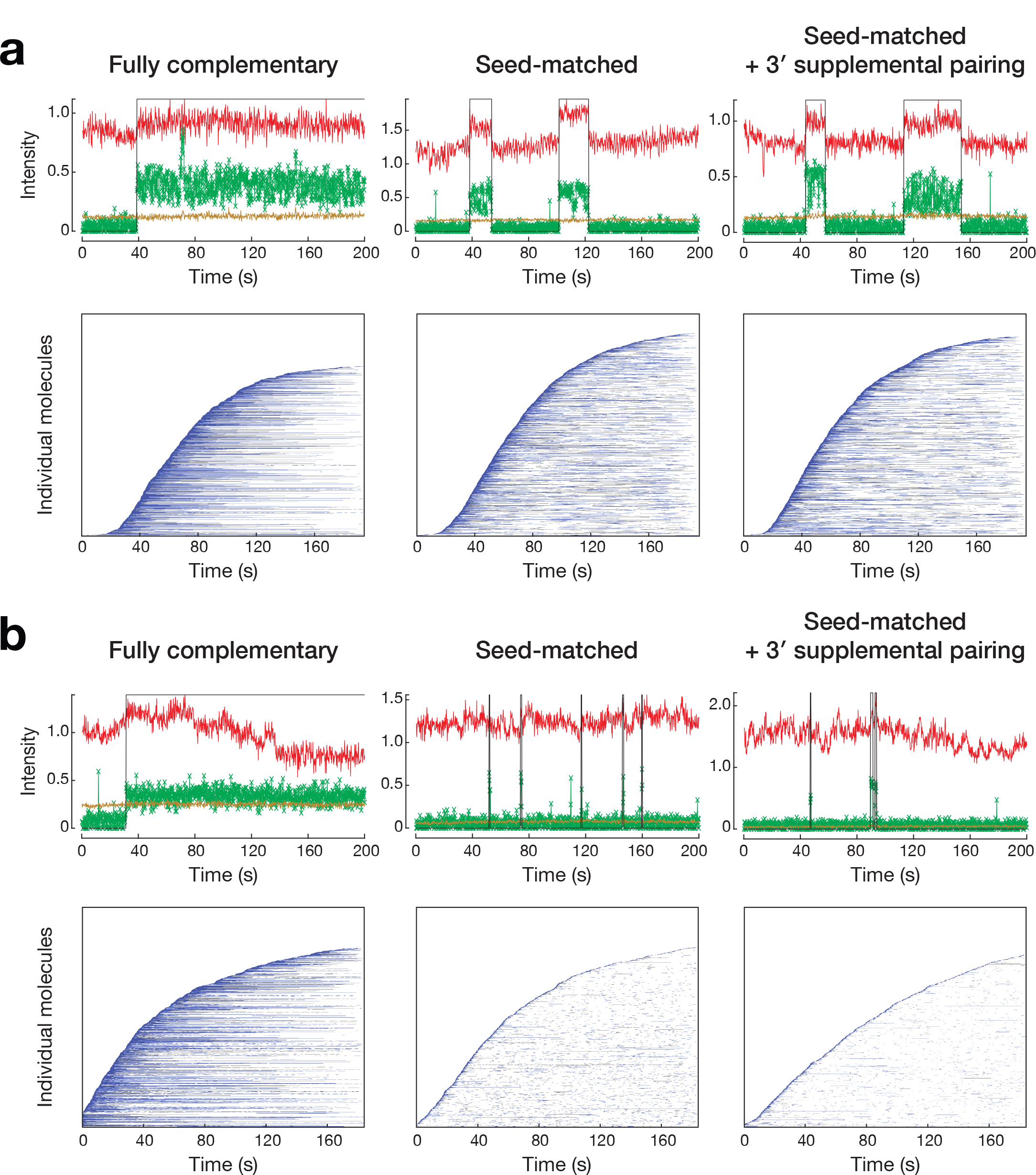
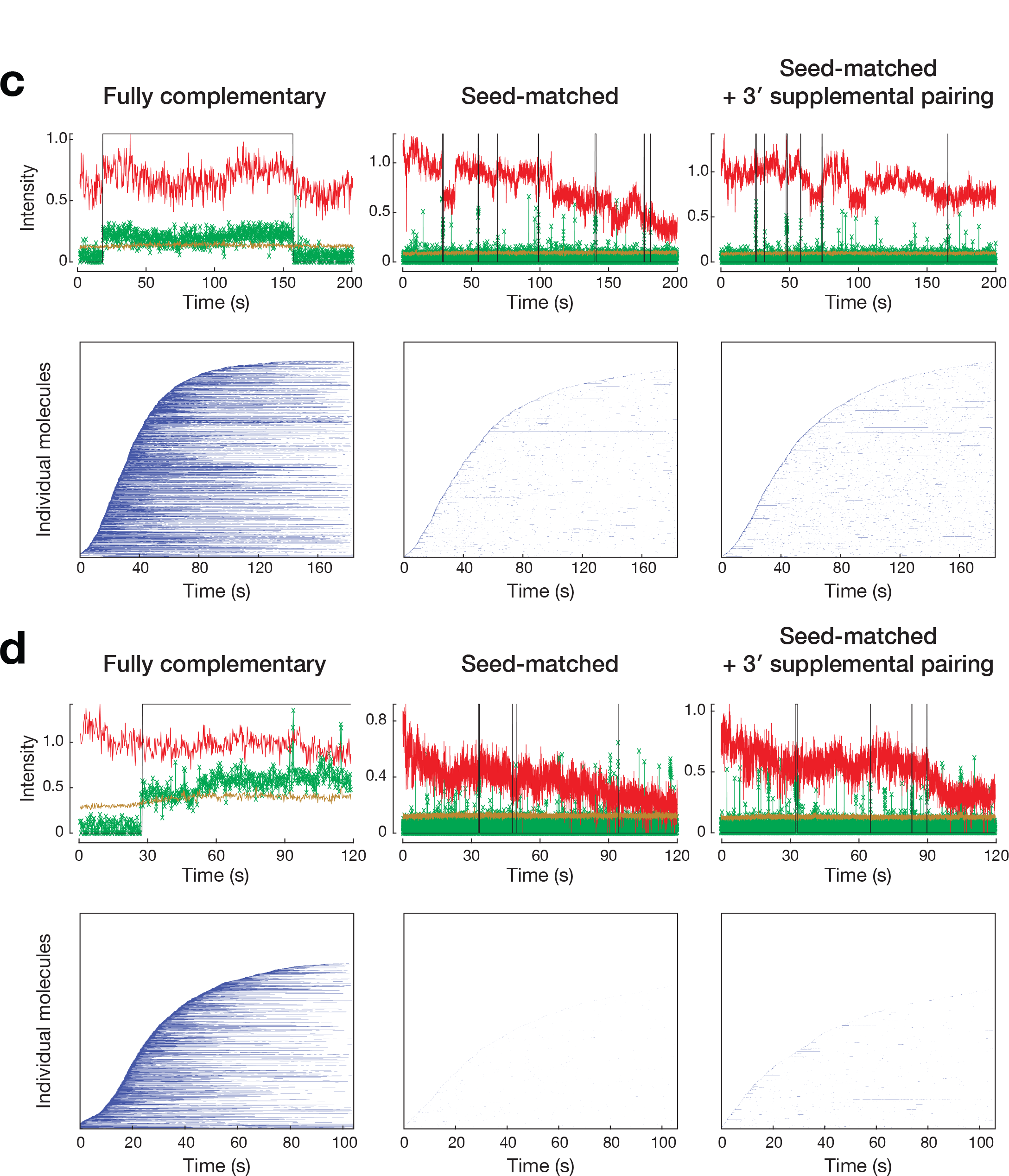
DNA-guided TtAgo Binding to and Departing from DNA Targets at Different Temperatures, Related to Figure 2. Representative fluorescence intensity time traces of DNA-guided TtAgo (green) binding DNA target (red) with different extents of complementarity. Light brown indicates background levels of green fluorescence, whereas the black line denotes binding events detected by the pipeline after event filtering (minimal duration and gap closing; Manual – Obtaining the binding traces). Fluorescence intensity is expressed in thousands of photons. Rastergrams summarize traces of individual target molecules, each in a single row and sorted according to their arrival time, for different guide:target pairings. Experiments were performed at 23°C (**a**), 37°C (**b**), 45°C (**c**) and 55°C (**d**).

**Figure S3.**
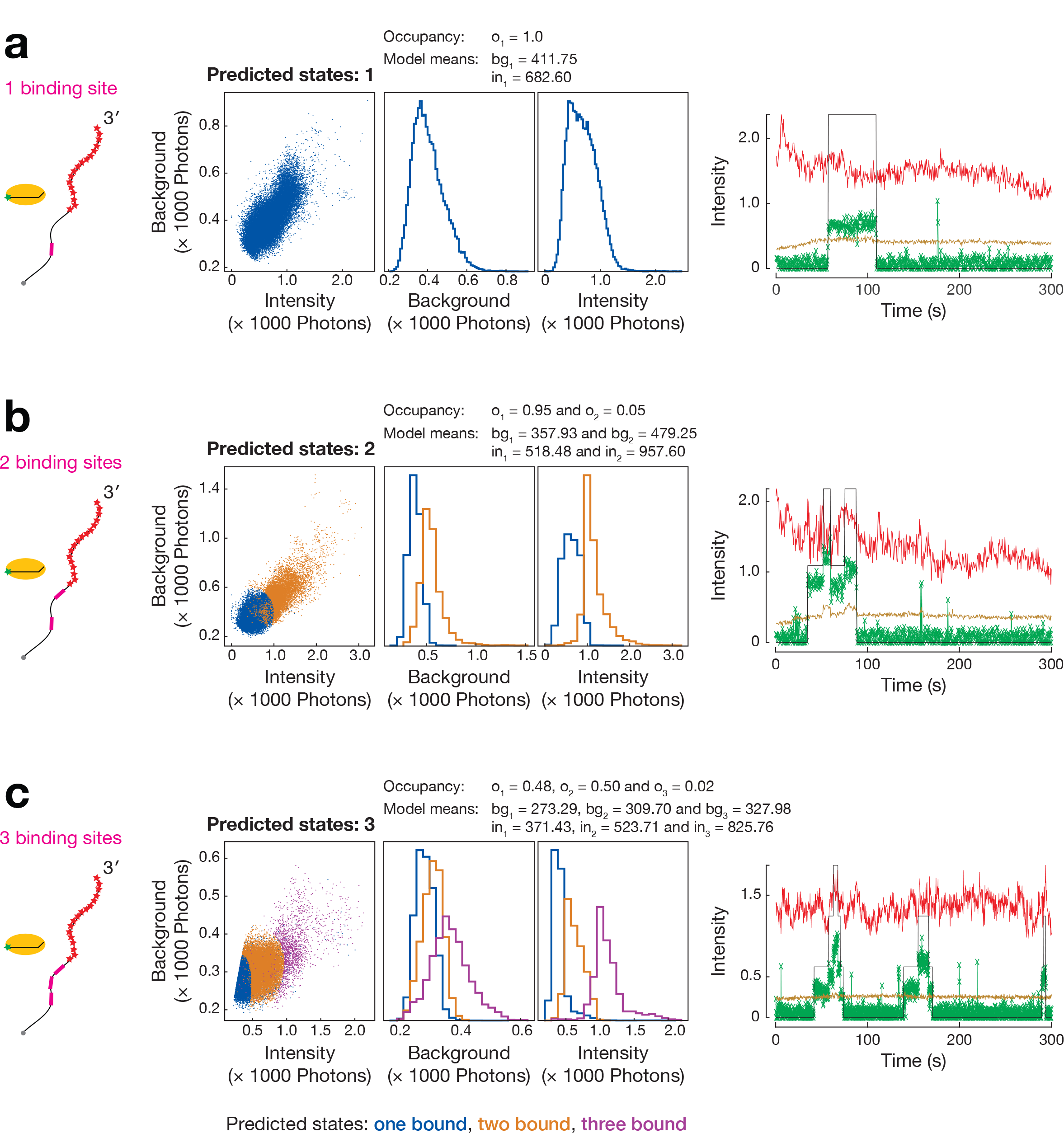
DNA-guided TtAgo Binding to Multiple, Fully Complementary Binding Sites and Prediction of the Number of States by VBEM-MGHMM, Related to Figure 3. Experimental setup to detect TtAgo:guide interactions with target DNA containing one (**a**), two (**b**) or three (**c**) binding site(s) complementary to the DNA guide. The number of states was estimated by VBEM-MGHMM. Occupancy, mean intensity of background and mean intensity of binding event are indicated for each state. Representative fluorescence intensity time traces of DNA-guided TtAgo (green) binding DNA target (red) are shown. Light brown indicates background levels of green fluorescence, whereas the black line denotes binding events detected by the pipeline after VBEM-MGHMM analysis. Fluorescence intensity is expressed in thousands of photons. o: occupancy, bg: background, in: intensity.

**Figure S4.**
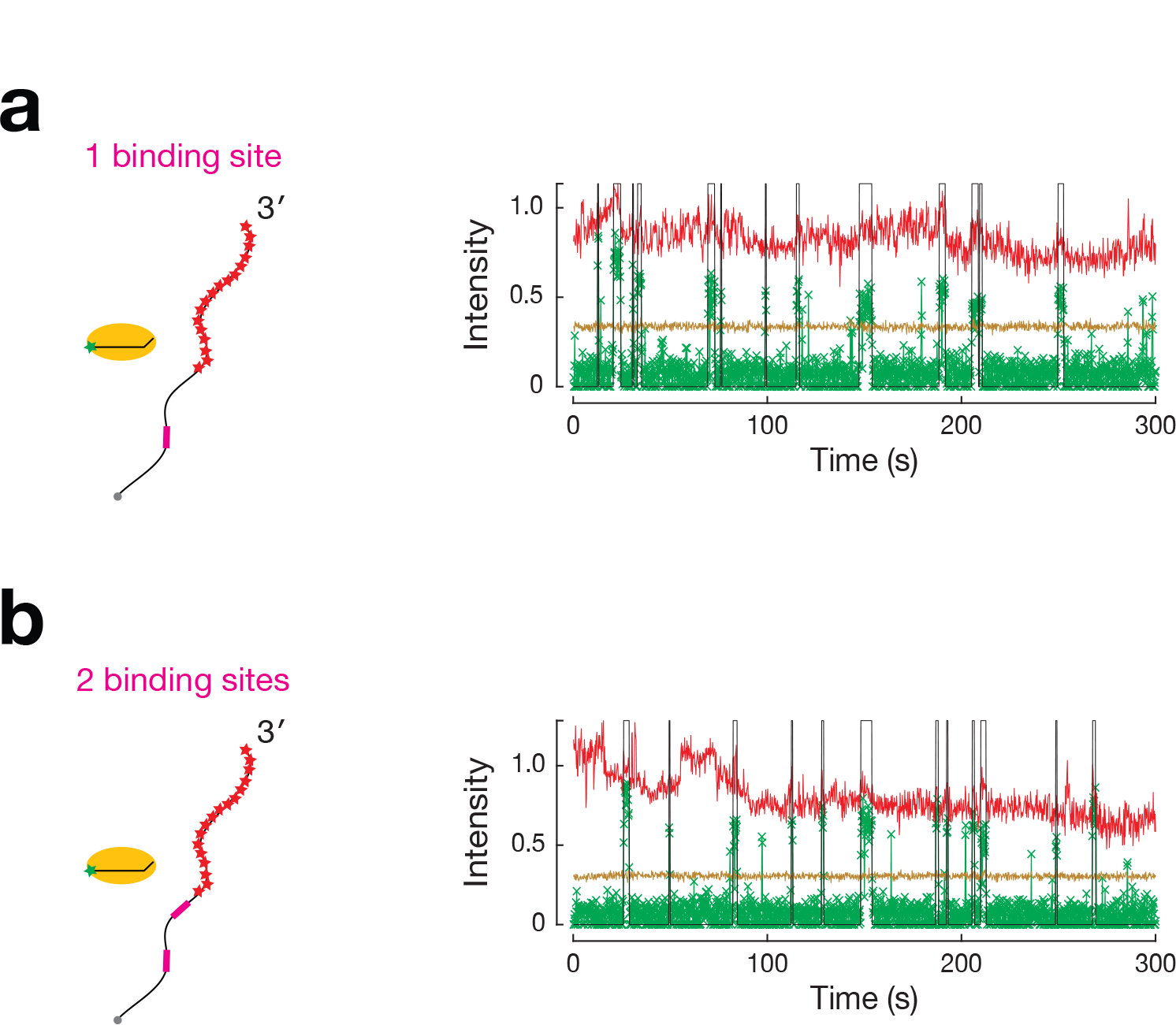
No Simultaneous Binding of TtAgo:Guide Complexes to a Target DNA with Two Seed-matched Binding Sites, Related to Figure 3. Experimental setup to measure TtAgo:guide complex interactions with target DNA containing one (**a**) or two (**b**) seed-matched binding site(s). Representative fluorescence intensity time traces of DNA-guided TtAgo (green) binding DNA target (red) are shown. Light brown indicates background levels of green fluorescence, whereas the black line denotes binding events detected by the pipeline after event filtering (minimal duration and gap closing; Manual – Obtaining the binding traces). Fluorescence intensity is expressed in thousands of photons.

**Figure S5.**
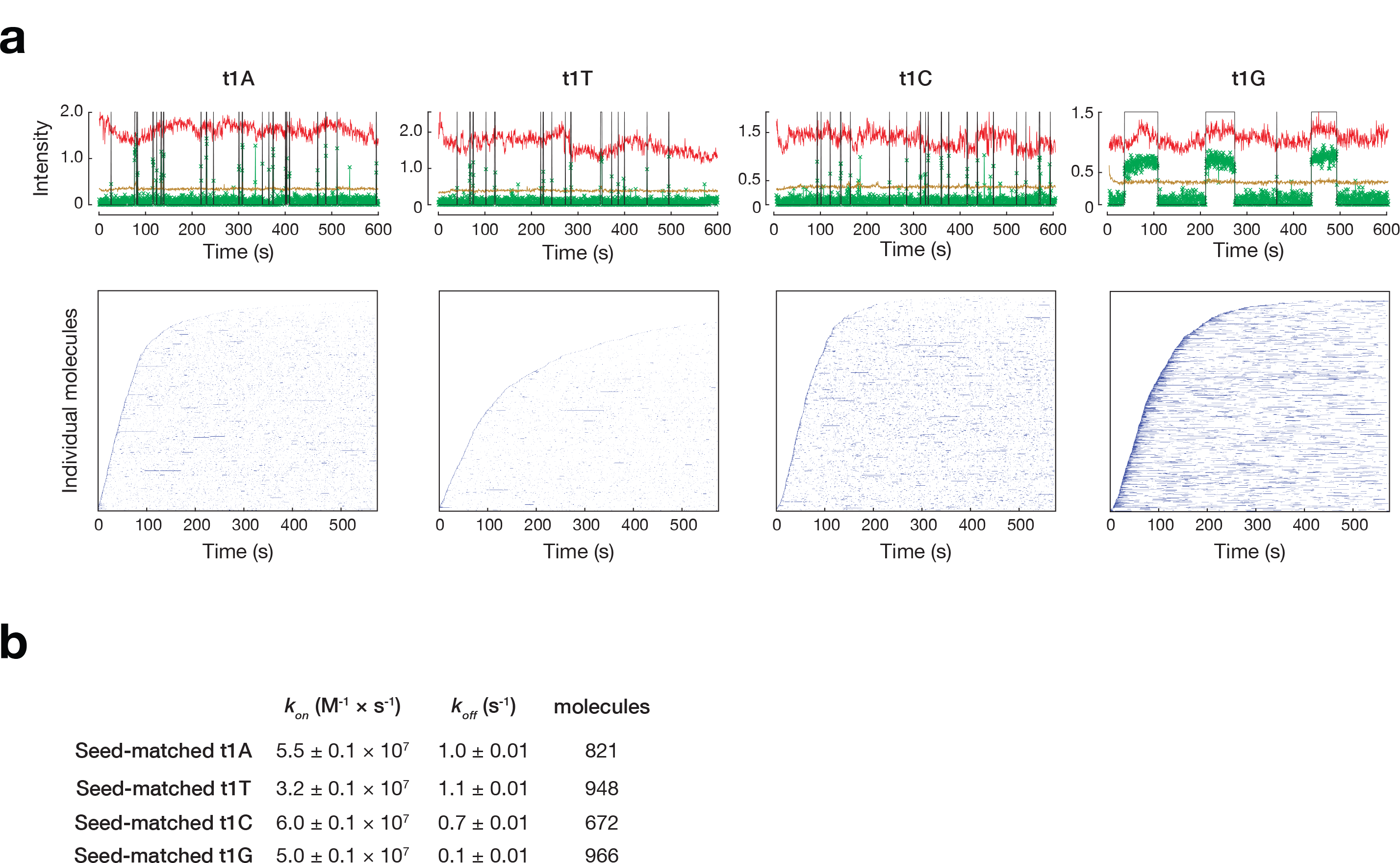
TtAgo:guide Complex Preferentially Binds to a t1G Target DNA, Related to Figure 3. (**a**) Representative fluorescence intensity time traces of DNA-guided TtAgo (green) binding DNA target (red) are shown. Light brown indicates background levels of green fluorescence, whereas the black line denotes binding events detected by the pipeline after event filtering (minimal duration and gap closing; Manual – Obtaining the binding traces). Fluorescence intensity is expressed in thousands of photons. Shown below are rastergram summaries of traces of individual target molecules, each in a single row and sorted according to their arrival time. (**b**) Comparison of *k*_*on*_ and *k*_*off*_ of DNA-guided TtAgo with different targets. Values were derived from data collected from several hundred individual DNA target molecules (indicated in the Table as number of molecules); error of bootstrapping is reported.

**Figure S6.**
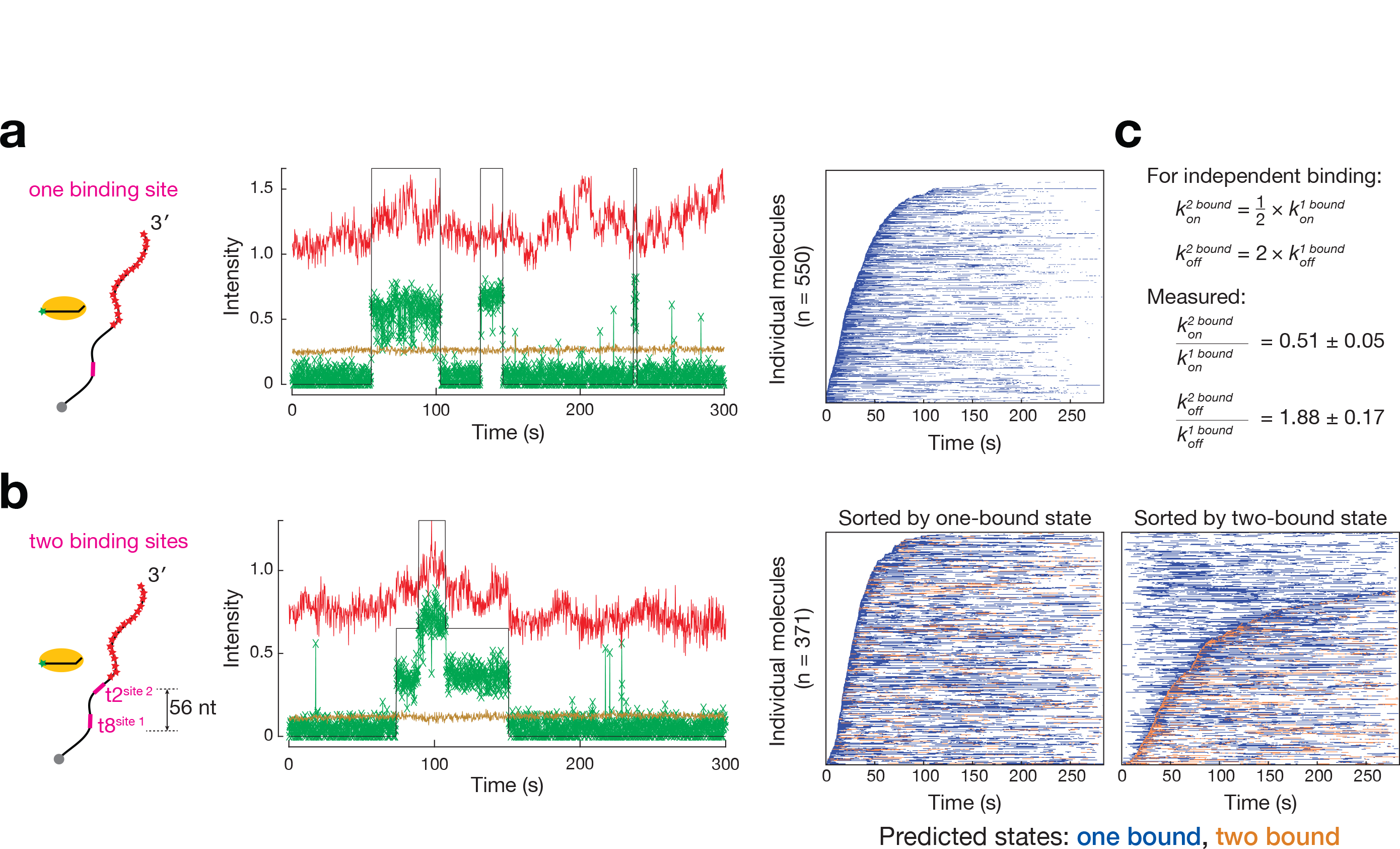
DNA-guided TtAgo Binds Independently to DNA Targets Containing Two Seed-Matched Sites with t1G, Relative to Figure 3. Representative fluorescence intensity time traces of DNA-guided TtAgo (green) binding DNA target (red) containing one binding site (**a**) or two binding sites spaced 56 nt apart from t8 to t2 (**b**). Light brown indicates background levels of green fluorescence, whereas the black line denotes binding events detected by the pipeline after VBEM-MGHMM analysis. Fluorescence intensity is expressed in thousands of photons. Representative rastergrams summarize traces of individual target molecules, each in a single row and sorted according to their arrival time. (**c**) Comparison of *k*_*on*_ and *k*_*off*_ of DNA-guided TtAgo with targets containing two or one binding site. Values are reported as mean ± standard deviation for three independent replicates.

**Table S1.**
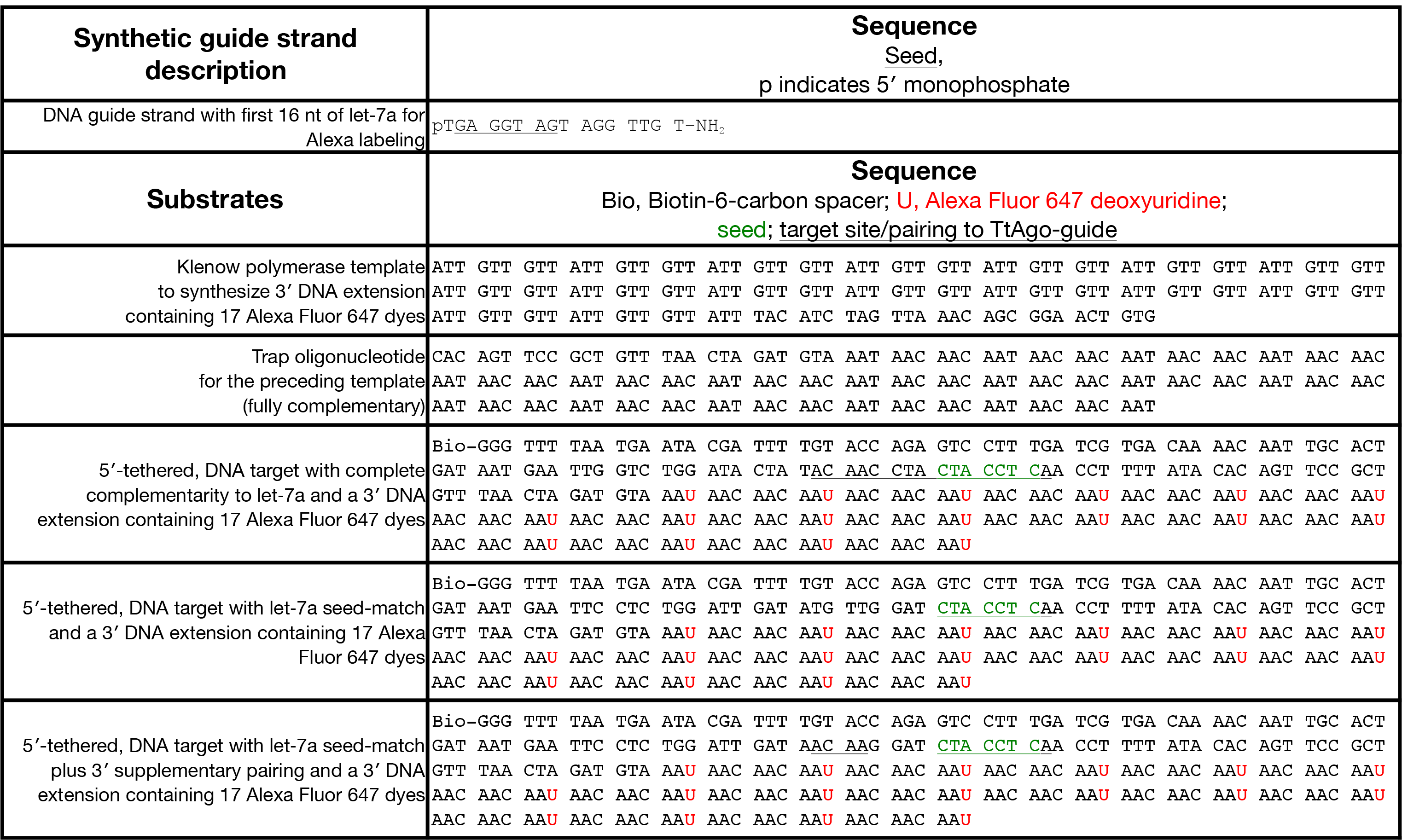
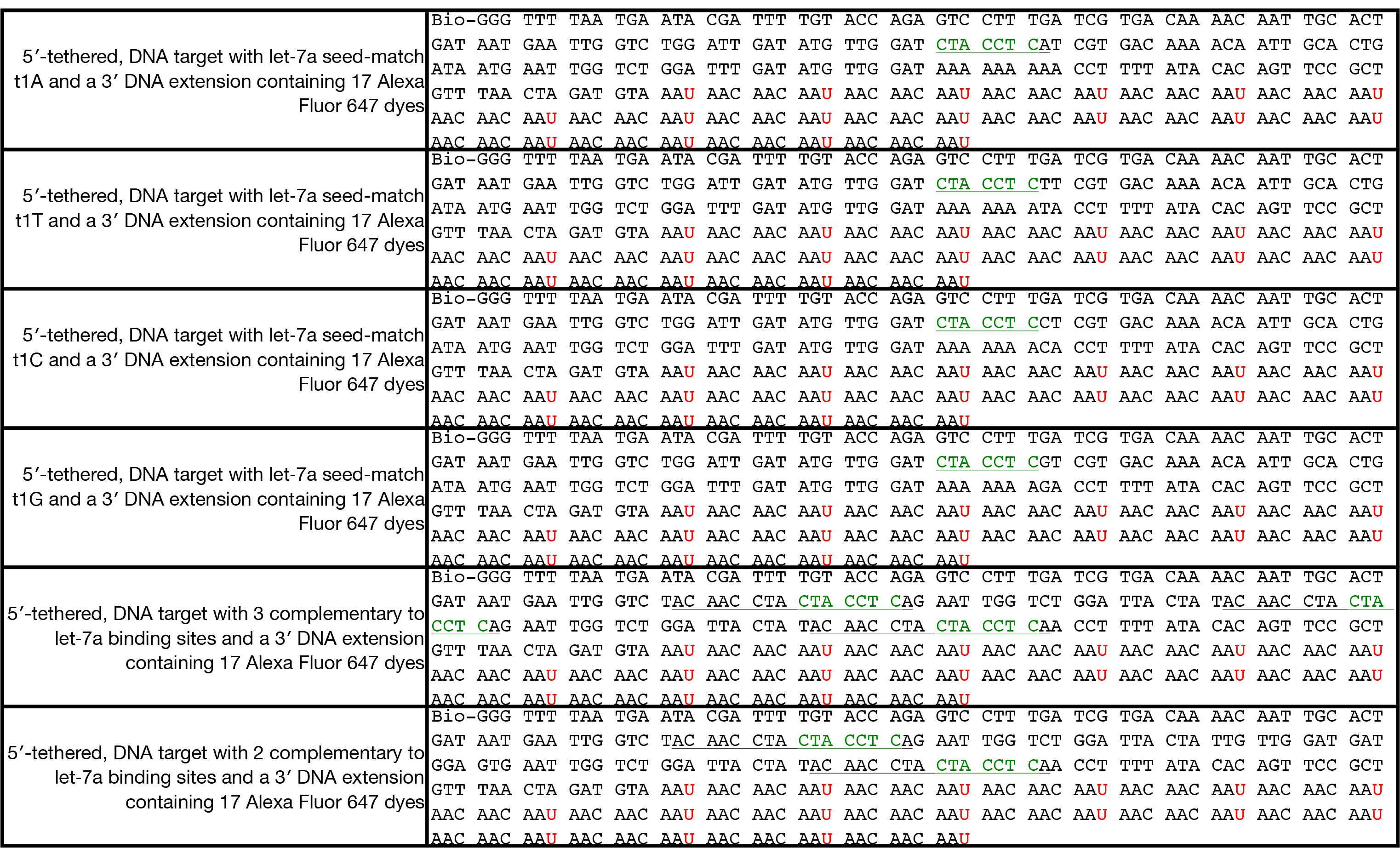
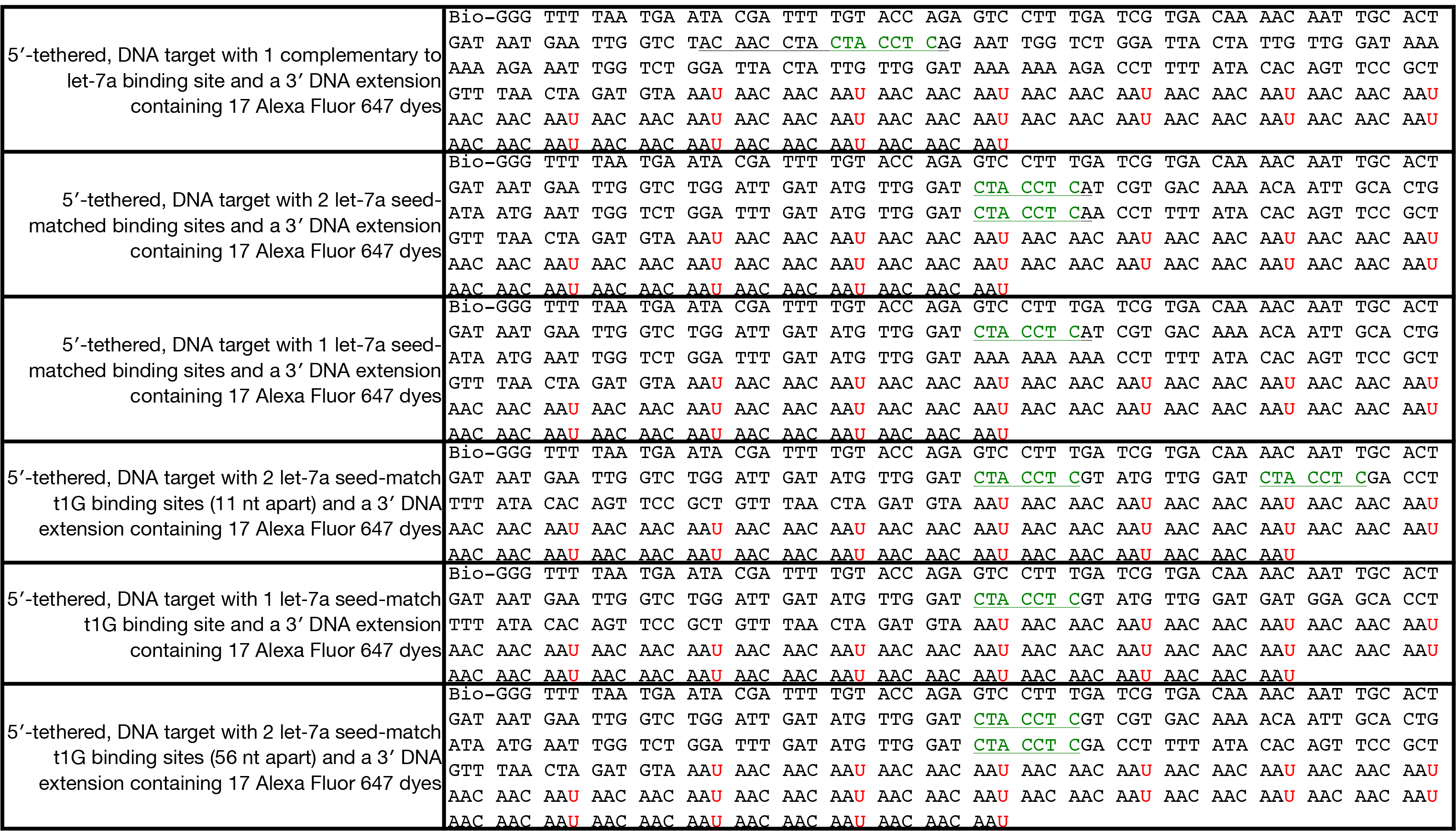
Related to Experimental Procedures. DNA oligos used in this study.

## References

1. Hoskins, A. A. et al. Ordered and dynamic assembly of single spliceosomes. Science 331, 1289–1295 (2011).

2. Friedman, L. J. & Gelles, J. Mechanism of transcription initiation at an activator-dependent promoter defined by single-molecule observation. Cell 148, 679–689 (2012).

3. Lee, H. W. et al. Real-time single-molecule co-immunoprecipitation analyses reveal cancer-specific Ras signalling dynamics. Nat Commun 4, 1505 (2013).

4. Chandradoss, S. D., Schirle, N. T., Szczepaniak, M., MacRae, I. J. & Joo, C. A Dynamic Search Process Underlies MicroRNA Targeting. Cell 162, 96–107 (2015).

5. Yao, C., Sasaki, H. M., Ueda, T., Tomari, Y. & Tadakuma, H. Single-Molecule Analysis of the Target Cleavage Reaction by the Drosophila RNAi Enzyme Complex. Mol Cell 59, 125–132 (2015).

6. Arauz, E., Aggarwal, V., Jain, A., Ha, T. & Chen, J. Single-Molecule Analysis of Lipid-Protein Interactions in Crude Cell Lysates. Anal Chem 88, 4269–4276 (2016).

7. Friedman, L. J. & Gelles, J. Multi-wavelength single-molecule fluorescence analysis of transcription mechanisms. Methods 86, 27–36 (2015).

8. Hansen, S. R., Rodgers, M. L. & Hoskins, A. A. in 581 (eds Spies, M. & Chemla, Y. R.) 83–104 (Academic Press, 2016).

9. Blanco, M. R. et al. Single Molecule Cluster Analysis dissects splicing pathway conformational dynamics. Nat Methods 12, 1077–1084 (2015).

10. van Vliet, L. J., Sudar, D. & Young, I. T. Digital Fluorescence Imaging Using Cooled CCD Array Cameras invisible. JE Celis (eds); Cell Biol. Second Eddition 3, 109–120 (1998).

11. Salomon, W. E., Jolly, S. M., Moore, M. J., Zamore, P. D. & Serebrov, V. Single-Molecule Imaging Reveals that Argonaute Reshapes the Binding Properties of Its Nucleic Acid Guides. Cell 162, 84–95 (2015).

12. Crocker, J. C. & Grier, D. G. Methods of Digital Video Microscopy for Colloidal Studies. Journal of Colloid and Interface Science 179, 298–310 (1996).

13. Smith, C. S. et al. Nuclear accessibility of β-actin mRNA is measured by 3D single-molecule real-time tracking. J Cell Biol 209, 609–619 (2015).

14. Shcherbakova, I. et al. Alternative spliceosome assembly pathways revealed by single-molecule fluorescence microscopy. Cell Rep 5, 151–165 (2013).

15. Hua, B. et al. The Single-Molecule Centroid Localization Algorithm Improves the Accuracy of Fluorescence Binding Assays. Biochemistry 57, 1572–1576 (2018).

16. Hui, J., Jiankun, Y. & Xiujian, L. Minimum variance unbiased subpixel centroid estimation of point image limited by photon shot noise. J. Opt. Soc. Am. A 27, 2038–2045 (2010).

17. Smith, C. S., Joseph, N., Rieger, B. & Lidke, K. A. Fast, single-molecule localization that achieves theoretically minimum uncertainty. Nat Methods 7, 373–375 (2010).

18. Bo, Z., Josiane, Z. & Jean-Christophe, O.-M. Gaussian approximations of fluorescence microscope point-spread function models. Appl. Opt. 46, 1819–1829 (2007).

19. Brown, C. M., Reilly, A. & Cole, R. W. A Quantitative Measure of Field Illumination. J Biomol Tech 26, 37–44 (2015).

20. Qin, F., Auerbach, A. & Sachs, F.A Direct Optimization Approach to Hidden Markov Modeling for Single Channel Kinetics. Biophysical Journal 79, 1915–1927 (2000).

21. Andrec, M., Levy, R. M. & Talaga, D. S. Direct Determination of Kinetic Rates from Single-Molecule Photon Arrival Trajectories Using Hidden Markov Models. The Journal of Physical Chemistry A J. Phys. Chem. A 107, 7454–7464 (2003).

22. McKinney, S. A., Joo, C. & Ha, T. Analysis of Single-Molecule FRET Trajectories Using Hidden Markov Modeling. Biophysical Journal 91, 1941–1951 (2006).

23. Low-Nam, S. T. et al. ErbB1 dimerization is promoted by domain co-confinement and stabilized by ligand binding. Nature Structural & Molecular Biology 18, 1244–1244 (2011).

24. Greenfeld, M., Pavlichin, D. S., Mabuchi, H. & Herschlag, D. Single Molecule Analysis Research Tool (SMART): an integrated approach for analyzing single molecule data. PLoS One 7, e30024 (2012).

25. Beal, M. J. Variational algorithms for approximate Bayesian inference 2003).

26. MacKay, D. J. C. Information theory, inference, and learning algorithms (Cambridge University Press, Cambridge, UK; New York, 2003).

27. Persson, F., Lindén, M., Unoson, C. & Elf, J. Extracting intracellular diffusive states and transition rates from single-molecule tracking data. Nat Methods 10, 265–269 (2013).

28. Bronson, J. E., Fei, J., Hofman, J. M., Gonzalez, R. L. & Wiggins, C. H. Learning Rates and States from Biophysical Time Series: A Bayesian Approach to Model Selection and Single-Molecule FRET Data. Biophysical Journal 97, 3196–3205 (2009).

29. van de Meent, J. W., Bronson, J. E., Wiggins, C. H. & Gonzalez, R. L. Empirical Bayes methods enable advanced population-level analyses of single-molecule FRET experiments. Biophys J 106, 1327–1337 (2014).

30. Monnier, N. et al. Inferring transient particle transport dynamics in live cells. Nat Methods 12, 838–840 (2015).

31. Bishop, C. M. Pattern Recognition and Machine Learning (Information Science and Statistics) (Springer-Verlag New York, Inc., Secaucus, NJ, USA, 2006).

32. Wang, Y., Sheng, G., Juranek, S., Tuschl, T. & Patel, D. J. Structure of the guide-strand-containing argonaute silencing complex. Nature 456, 209–213 (2008).

33. Swarts, D. C. et al. DNA-guided DNA interference by a prokaryotic Argonaute. Nature 507, 258–261 (2014).

34. Wang, Y. et al. Structure of an argonaute silencing complex with a seed-containing guide DNA and target RNA duplex. Nature 456, 921–926 (2008).

35. Wang, Y. et al. Nucleation, propagation and cleavage of target RNAs in Ago silencing complexes. Nature 461, 754–761 (2009).

36. Sheng, G. et al. Structure-based cleavage mechanism of Thermus thermophilus Argonaute DNA guide strand-mediated DNA target cleavage. Proc Natl Acad Sci U S A 111, 652–657 (2014).

37. Swarts, D. C. et al. Autonomous Generation and Loading of DNA Guides by Bacterial Argonaute. Mol Cell 65, 985–998.e6 (2017).

38. Jung, S. R. et al. Dynamic anchoring of the 3’-end of the guide strand controls the target dissociation of Argonaute-guide complex. J Am Chem Soc 135, 16865–16871 (2013).

39. Oshima, T. & Imahori, K. Description of Thermus thermophilus (Yoshida and Oshima) comb. nov., a Nonsporulating Thermophilic Bacterium from a Japanese Thermal Spa. International Journal of Systematic and Evolutionary Microbiology 24, 102–112 (1974).

40. Jo, M. H. et al. Human Argonaute 2 Has Diverse Reaction Pathways on Target RNAs. Mol Cell 59, 117–124 (2015).

41. Grimson, A. et al. MicroRNA targeting specificity in mammals: determinants beyond seed pairing. Mol Cell 27, 91–105 (2007).

42. Broderick, J. A., Salomon, W. E., Ryder, S. P., Aronin, N. & Zamore, P. D. Argonaute protein identity and pairing geometry determine cooperativity in mammalian RNA silencing. RNA 17, 1858–1869 (2011).

43. Wang, W. et al. The initial uridine of primary piRNAs does not create the tenth adenine that Is the hallmark of secondary piRNAs. Mol Cell 56, 708–716 (2014).

44. Schirle, N. T., Sheu-Gruttadauria, J., Chandradoss, S. D., Joo, C. & MacRae, I. J. Water-mediated recognition of t1-adenosine anchors Argonaute2 to microRNA targets. eLife 4, e07646 (2015).

45. Crawford, D. J., Hoskins, A. A., Friedman, L. J., Gelles, J. & Moore, M. J. Visualizing the splicing of single pre-mRNA molecules in whole cell extract. RNA 14, 170–179 (2008).

46. Aitken, C. E., Marshall, R. A. & Puglisi, J. D. An Oxygen Scavenging System for Improvement of Dye Stability in Single-Molecule Fluorescence Experiments. Biophysical Journal 94, 1826–1835 (2008).

47. Dave, R., Terry, D. S., Munro, J. B. & Blanchard, S. C. Mitigating unwanted photophysical processes for improved single-molecule fluorescence imaging. Biophys J 96, 2371–2381 (2009).

48. LaMont, C. H. & Wiggins, P. A. The Lindley paradox: The loss of resolution in Bayesian inference. arXiv:1610.09433v2 [math.ST] (2017).

